# Highly Contiguous Genome Assembly of *Drosophila prolongata* - a Model for Evolution of Sexual Dimorphism and Male-specific Innovations

**DOI:** 10.1101/2024.01.29.577853

**Authors:** David Luecke, Yige Luo, Halina Krzystek, Corbin Jones, Artyom Kopp

**Affiliations:** Department of Evolution and Ecology, University of California Davis, One Shields Ave Davis CA 95616; Biology Department of the University of North Carolina (UNC), 3159 Genome Sciences Building. 250 Bell Tower Drive. Chapel Hill, NC 27599

**Author notes:** Corresponding authors: David Luecke, Artyom Kopp.

**Keywords:** Genome, Drosophila, Sex Dimorphism

## Abstract

*Drosophila prolongata* is a member of the *melanogaster* species group and *rhopaloa* subgroup native to the subtropical highlands of southeast Asia. This species exhibits an array of recently evolved male-specific morphological, physiological, and behavioral traits that distinguish it from its closest relatives, making it an attractive model for studying the evolution of sexual dimorphism and testing theories of sexual selection. The lack of genomic resources has impeded the dissection of the molecular basis of sex-specific development and behavior in this species. To address this, we assembled the genome of *D. prolongata* using long-read sequencing and Hi-C scaffolding, resulting in a highly complete and contiguous (scaffold N50 2.2Mb) genome assembly of 220Mb. The repetitive content of the genome is 24.6%, the plurality of which are LTR retrotransposons (33.2%). Annotations based on RNA-seq data and homology to related species revealed a total of 19,330 genes, of which 16,170 are protein-coding. The assembly includes 98.5% of Diptera BUSCO genes, including 93.8% present as a single copy. Despite some likely regional duplications, the completeness of this genome suggests that it can be readily used for gene expression, GWAS, and other genomic analyses.

## Introduction

*Drosophila prolongata* is a member of the *melanogaster* species group and *rhopaloa* subgroup native to southeast Asia (Singh and Gupta 1977; Toda 1991). The species has a suite of recently evolved male-specific morphological traits (Figure 1), including increased foreleg size, leg pigmentation, wing pigmentation, reversed sexual size dimorphism, and an expanded number of leg chemosensory organs (Luecke, Rice, and Kopp 2022; Luecke and Kopp 2019; Luo et al. 2019). These traits are associated with derived behaviors, including male-male grappling and male leg vibration courtship displays, along with increased sexual dimorphism in cuticular hydrocarbon profiles (Amino and Matsuo 2023b; 2023a; Kudo et al. 2015; 2017; Luo et al. 2019; Setoguchi et al. 2014; Takau and Matsuo 2022; Toyoshima and Matsuo 2023).

**Figure 1.**
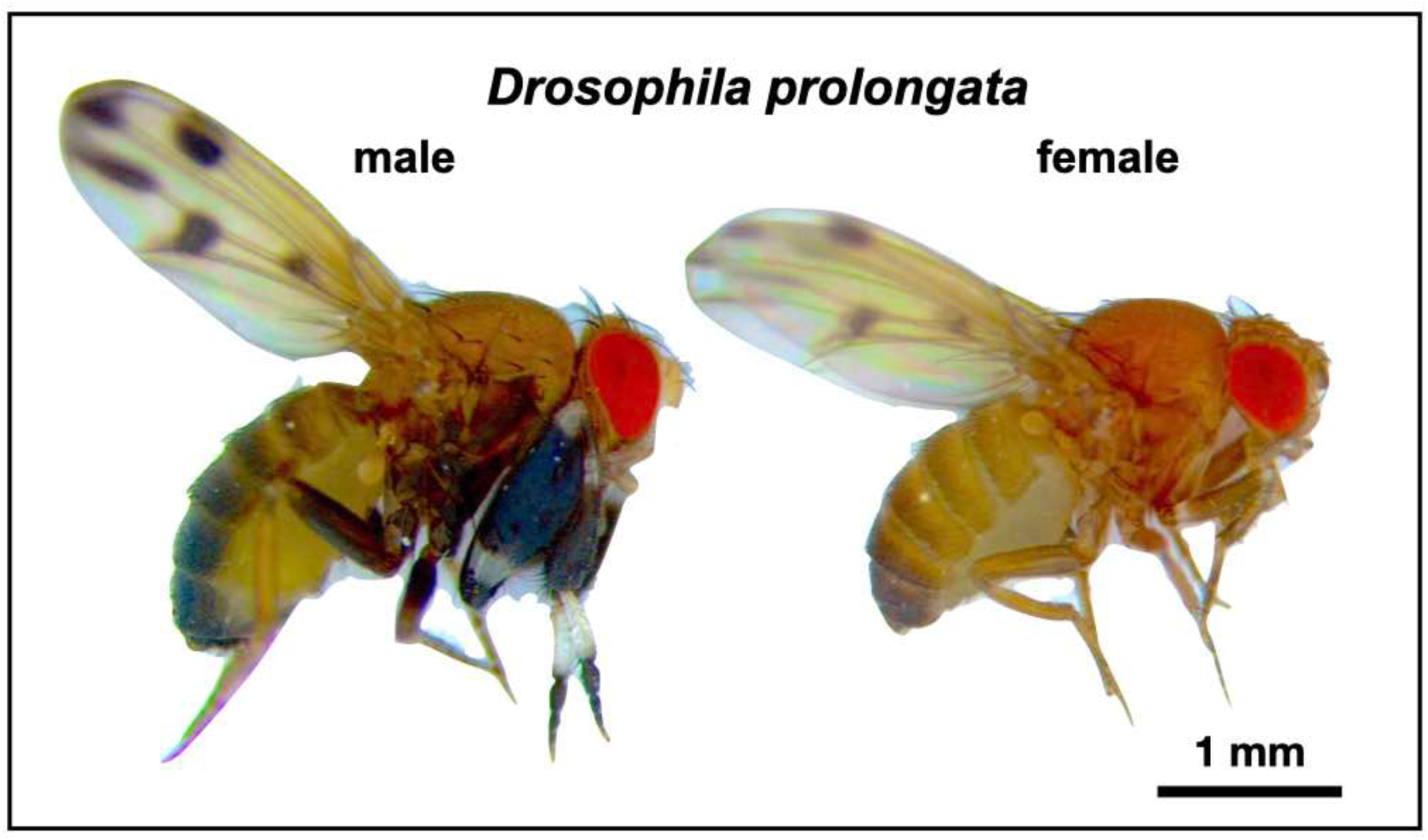
*Drosophila prolongata* has a suite of recently evolved male-specific traits, ideal for studying the evolution of sexual dimorphism. Most noticeable is the size and pigmentation banding of front legs in males. Other sexually dimorphic characteristics include wing spots, eye shape, pigmentation, and increased length of second and third legs.

The phylogenetic proximity to the model species *D. melanogaster* and available genome sequences for closely related species *D. rhopaloa* and *D. carrolli* (Kim et al. 2021), which lack these derived traits, make this species a promising system to study the genetics of sexually dimorphic development, physiology, and behavior. A reference genome assembly and annotation for *D. prolongata* benefits such work as it would provide insight into the genomic evolutionary patterns associated with the evolution of the novel traits in *D. prolongata*. Presented here is a highly complete and contiguous assembly based on long-read Pacific Biosciences sequencing and Hi-C scaffolding, along with annotations for both *D. prolongata* and *D. carrolli* using *D. melanogaster* sequence homology and gene models based on RNA sequencing evidence and ab initio predictions.

## Materials and Methods

### Genome line generation

The isofemale SaPa01 line and BaVi44 line were collected in SaPa and BaVi, Vietnam, respectively, by Dr. Hisaki Takamori in September 2004. Virgin females were collected by isolating adults within four hours of emergence. Four generations of full-sib matings were carried out to produce the genomic strain SaPa_ori_Rep25-2-1-1 (“Sapa_PacBio”). Fly strains were maintained at room temperature on standard cornmeal food provided by the UC Davis Fly Kitchen with filter paper for environment structure and pupariation substrate.

### Tissue collection

For genome assembly/scaffolding, adult male flies from the genome strain were moved onto nutrient-free agar media for at least one day to reduce microbial load, then collected into 1.5mL tubes and flash-frozen in liquid nitrogen. Fifty frozen adult male individuals were sent on dry ice to Dovetail Genomics (Cantata Bio. LLC, dovetailgenomics.com) for DNA extraction, sequencing, and assembly. For gene expression data used in annotation, whole forelegs were dissected from carbon dioxide anesthetized males and females of the SaPa01 isofemale line, along with dissected heads from each sex of the genome strain.

### Sequencing and assembly

All genomic DNA extraction, sequencing, and assembly were carried out by Dovetail Genomics (Cantata Bio LLC, Scotts Valley, CA, USA). Genomic DNA was extracted with the Qiagen HMW genomic extraction kit (Qiagen, Germantown, MD, USA). DNA samples were quantified using a Qubit 2.0 Fluorometer (Life Technologies, Carlsbad, CA, USA). The PacBio SMRTbell library (∼20kb) for PacBio Sequel was constructed using SMRTbell Express Template Prep Kit 2.0 (PacBio, Menlo Park, CA, USA) using the manufacturer-recommended protocol. The library was bound to polymerase using the Sequel II Binding Kit 2.0 (PacBio) and loaded onto PacBio Sequel II. Sequencing was performed on PacBio Sequel II 8M SMRT cells, generating 16 gigabases of data. An initial assembly based on 1.2M PacBio reads was produced using FALCON (Chin et al. 2016) with Arrow polishing.

A Dovetail HiC library was prepared similarly as described previously (Lieberman-Aiden et al. 2009). Briefly, for each library, chromatin was fixed in place with formaldehyde in the nucleus and then extracted. Fixed chromatin was digested with DpnII, the 5’ overhangs filled in with biotinylated nucleotides, and free blunt ends were ligated subsequently. After ligation, crosslinks were reversed, and the DNA was purified from protein. Purified DNA was treated to remove biotin that was not internal to ligated fragments. The DNA was then sheared to ∼350 bp mean fragment size, and sequencing libraries were generated using NEBNext Ultra enzymes and Illumina-compatible adapters. Biotin-containing fragments were isolated using streptavidin beads before PCR enrichment of each library. The libraries were sequenced on an Illumina HiSeq X to a target depth of 30x coverage.

The input *de novo* assembly and Dovetail HiC library reads were used as input data for HiRise, a software pipeline designed specifically for using proximity ligation data to scaffold genome assemblies (Putnam et al. 2016). Dovetail HiC library sequences were aligned to the draft input assembly using a modified SNAP read mapper (http://snap.cs.berkeley.edu). The separations of Dovetail HiC read pairs mapped within draft scaffolds were analyzed by HiRise to produce a likelihood model for genomic distance between read pairs, and the model was used to identify and break putative misjoins, to score prospective joins, and make joins above a threshold. A second HiRise assembly was generated with additional HiC sequencing and the HiRise software pipeline.

RNA was extracted using TRIzol (Invitrogen, Waltham, MA, USA). For foreleg RNA, multiplexed stranded cDNA sequencing libraries were prepared using the NEBNext Ultra Directional RNA Library Prep Kit for Illumina (New England BioLabs, Ipswich, MA, USA) using poly(A) isolation magnetic beads. Libraries were sequenced on the Illumina HiSeq4000 platform by the UC Davis Genome Center. For head RNA, cDNA sequencing libraries were constructed using the TruSeq Stranded RNA Kit (Illumina, San Diego, CA) and sequenced on the Illumina HiSeq4000 platform by Novogene (https://www.novogene.com/us-en/). Raw RNA-seq reads and assembled genome can be accessed with NCBI BioProject PRJNA1057277. Transcripts were assembled using Trinity 2.4.0 (Haas et al. 2013) with default options for stranded data.

### Gene prediction and annotation

Homology-based annotations were generated using Liftoff 1.5.1 (Shumate and Salzberg 2021) with minimap2 2.17 (Li 2018) alignment based on the *D. melanogaster* GCF000001215.4 release 6 (Hoskins et al. 2015) *D. elegans* GCF000224195.1 2.0, and *D. rhopaloa* GCF000236305.1 2.0 (Kim et al. 2021) annotations downloaded from FlyBase (Gramates et al. 2022). Liftoff was run with the copies option and percent identity 0.80. Additional gene models were inferred using MAKER 3.01.02 (Holt and Yandell 2011) with BLAST 2.11.0 (Camacho et al. 2009) and repeat masker 4.0.7, using EST evidence from the Trinity transcripts assembled based on foreleg and head RNA and protein homology evidence based on the combined protein sets from the *D. melanogaster* and *D. elegans* annotations also used for Liftoff. The annotations from different sources were then combined using gffcompare 10.4 (Pertea and Pertea 2020), genometools 1.5.9 (Gremme, Steinbiss, and Kurtz 2013), and custom Python 3.7.6 scripts available at https://github.com/dluecke/annotation_tools.

### Removal of duplicate scaffolds

BUSCO (Manni et al. 2021) analysis of the Dovetail HiRise using the diptera_ocb10 lineage dataset revealed 200 complete but duplicated benchmark genes (Table S1), indicating potential duplicated regions in the assembly. Scaffolds were assessed for BUSCO benchmark gene content and sorted by the percentage of duplicated BUSCO genes. 53 candidate scaffolds, ranging from 20,819bp to 39,990,007bp, contained at least one duplicated benchmark BUSCO gene (Table S1). Inspection of MUMmer (Marçais et al. 2018) alignments between duplicate-containing candidates and scaffolds with alternate copies of the duplicated benchmark genes showed complete alignment across 27 of the candidate scaffolds (Figure S1). These 27 scaffolds (ranging from 20,819bp to 541,551bp) were considered fully duplicated and split from the assembly and annotation (Files S1, S2, and S3) using SAMtools 1.15.1 (Li et al. 2009). Custom Python pandas 1.1.2 (McKinney 2010), and R 4.0.3 (R Core Team 2020, https://www.R-project.org) for scaffold sorting by BUSCO scores, splitting assembly and annotation, and inspecting genome alignments are available at https://github.com/dluecke/annotation_tools.

### Identification of duplicate genes

The remaining duplicated genes in the *D. prolongata* deduplicated annotation were identified using reciprocal BLAST. Strand oriented regions corresponding to all “gene” features in both *D. prolonogata* and *D. rhopaloa* annotations were extracted from their respective assemblies using bedtools 2.29.2 (Quinlan and Hall 2010). *D. prolongata* gene regions were searched against all *D. prolongata* and all *D. rhopaloa* gene regions using blastn 2.14.1 (Camacho et al. 2009). BLAST results were combined and sorted by match alignment bit score, then duplicate status was assigned to pairs of *D. prolongata* genes if both regions had higher match scores with the corresponding *D. prolongata* region than to any gene region from *D rhopaloa*. Custom Bash and Python scripts used in this process are available at https://github.com/dluecke/annotation_tools.

### Repeat analysis

Tandem repeats were annotated with Tandem Repeat Finder 4.09.1 (Benson 1999). A *de novo* library of classified repetitive element models was created using RepeatModeler 2.0 (Flynn et al. 2020). To reduce the run-to-run variations, repeat classification was based on five independent RepeatModeler runs with the following random seeds: 1681089287, 1687990919, 1683413925, 1683532158, and 1683532058. Custom R and Bash scripts are available at https://github.com/yige-luo/Repeat_analysis.

### Assembly and annotation evaluation

Assembly contiguity statistics were provided by Dovetail. Reference annotations *D. melanogaster* GCF_000001215.4 and *D. rhopaloa* GCF_018152115.1 were downloaded from the NCBI genomes database. Assembly completeness was assessed with BUSCO 5.3.2 (Manni et al. 2021) using the diptera_ocb10 lineage dataset, HMMER 3.1b2, and Mmseqs 5.34c21f2. Whole genome alignment between *D. prolongata* and *D. rhopaloa* assemblies was performed with MUMmer 4.0.0 (Marçais et al. 2018) using nucmer alignment with a minimum exact match 1000bp for alignment with *D. rhopaloa* and 500bp for *D. melanogaster* alignment, and mummerplot plus custom Bash and R scripts (https://github.com/dluecke/annotation_tools) for visualization. Annotation statistics were found with genometools 1.5.9 (Gremme, Steinbiss, and Kurtz 2013). Transcripts were extracted from annotations using gffread 0.9.12 (Pertea and Pertea 2020), and transcript completeness was assessed using the transcriptome mode of BUSCO.

## Results and Discussion

### Assembly contiguity

The Dovetail HiRise assembly scaffolding method (Figure 2) produced an assembly for *D. prolongata* with higher contiguity than the existing *D. rhopaloa* and *D. carrolli* assemblies, approaching the contiguity of the latest *D. melanogaster* reference (Table 1) as measured by N50. Whole genome alignments of the *D. prolongata* assembly to *D. rhopaloa* and *D. melanogaster* references (Figure 3A) show long stretches of high identity with *D. rhopaloa* spanning nearly all large scaffolds.

**Figure 2.**
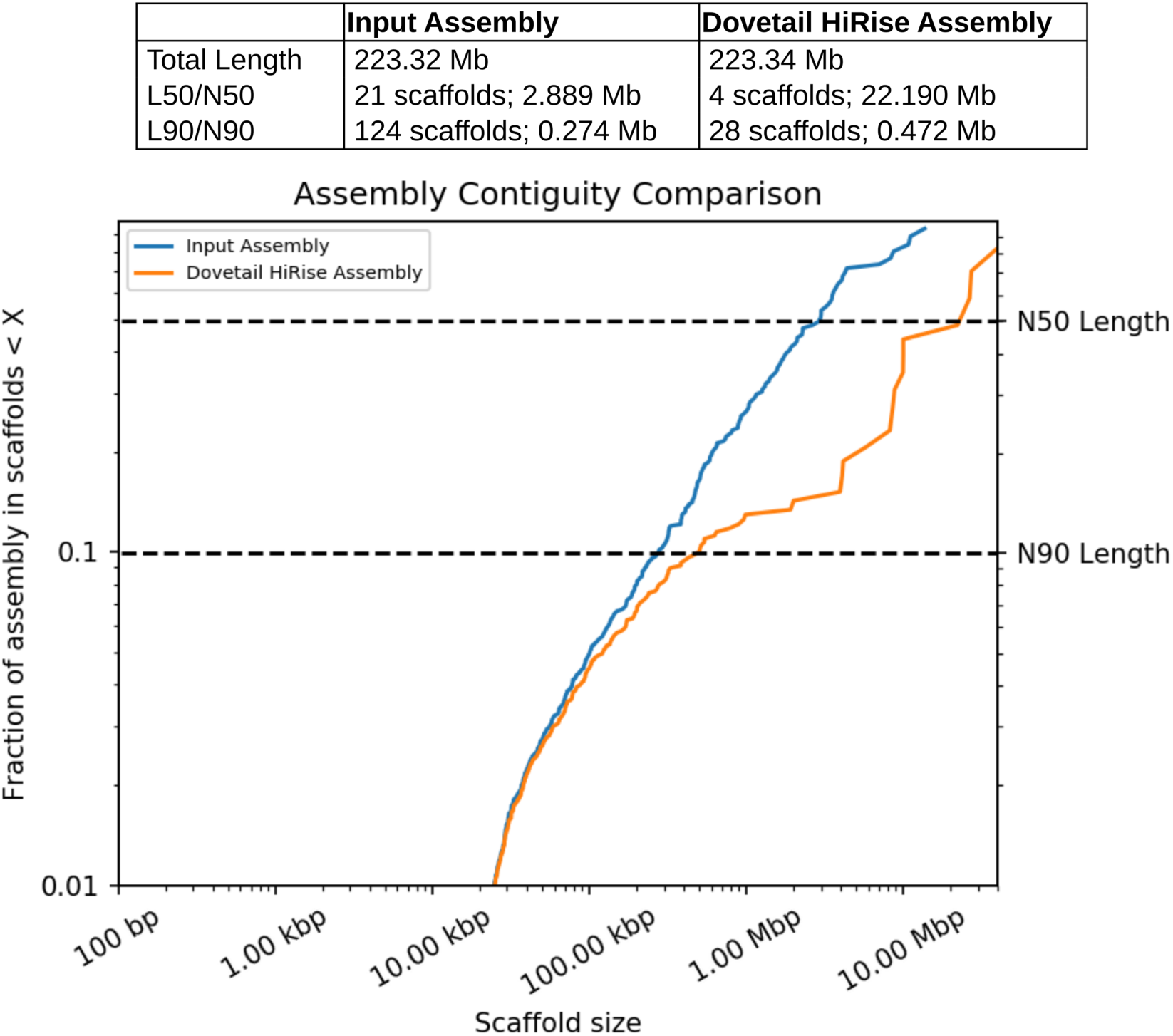
Dovetail assembly process generates high contiguity assembly. Comparison between initial PacBio FALCON with Arrow polished assembly (“Input Assembly”) and final assembly generated by Dovetail HiC scaffolding method (“HiRise Assembly”), provided by Dovetail genomics. Each curve shows the fraction of the total length of the assembly in scaffolds of a given length or smaller. Scaffolds shorter than 1kb are excluded.

**Figure 3.**
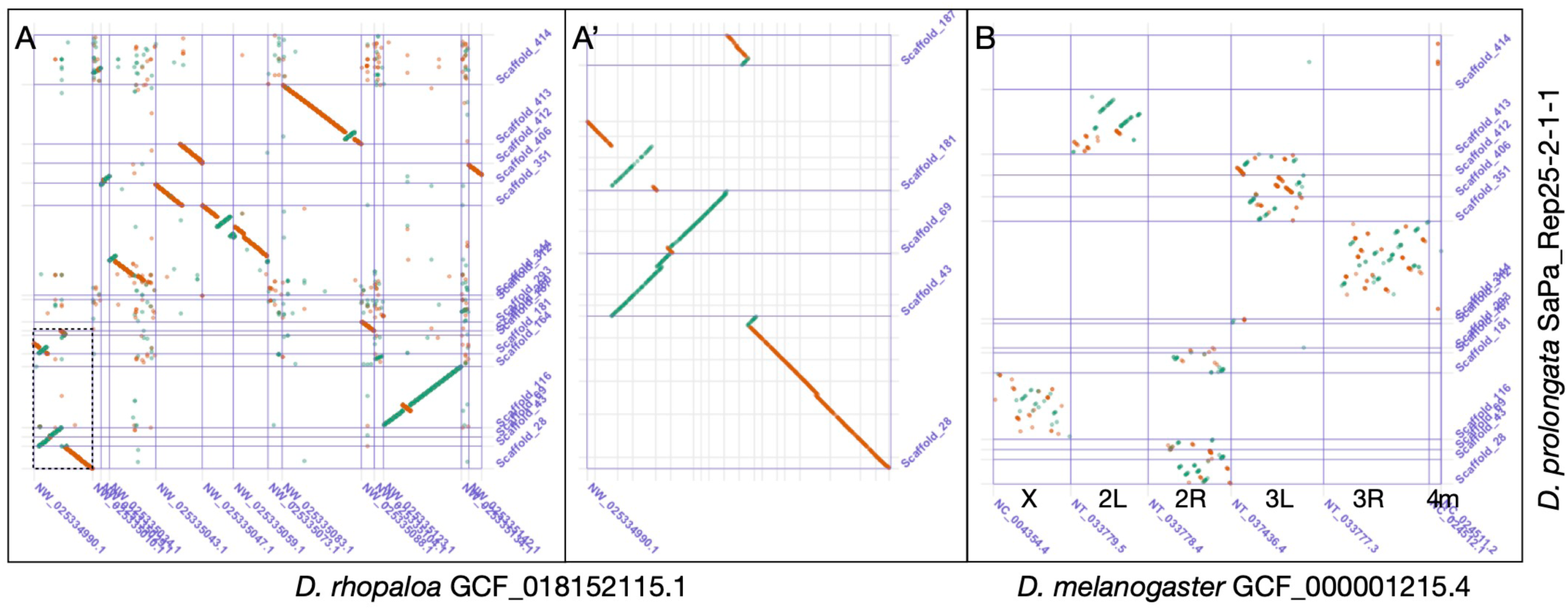
Whole genome alignments between major scaffolds of *D. prolongata* (>1Mb) assembly and reference assemblies. Sense matches are shown in green, and anti-sense matches in orange. (A) Alignment to *D. rhopaloa* reference based on minimum 1000bp matches, showing reference scaffolds >2.5Mb as ordered in assembly; boxed area is expanded in panel A’. (A’) Zoom on portion of alignment A, showing regional duplication and inversion. (B) Alignment to major chromosome arms from *D. melanogaster* assembly, based on minimum 500bp matches. Large stretches of contiguity with limited large inversions are evident between *D. prolongata* and *D. rhopaloa* (A), while conservation of each chromosome arm’s content along with considerable intra-arm rearrangement is seen between *D. prolongata* and *D. melanogaster* (B). A duplication spanning an inversion is evident between Scaffold_43 and Scaffold_181 (A’).

**Table 1:** Statistics for assembly contiguity and completeness of *D. prolongata* assembly alongside previously published *D. carrolli* GCA_018152295.1 assembly (Kim et al. 2021), reference assemblies *D. rhopaloa* GCF_018152115.1 and *D. melanogaster* GCF_000001215.4. BUSCO statistics are for the 3285 genes in the diptera_odb10 benchmark set.

| Assembly | *D. prolongata* | *D. carrolli* | *D. rhopaloa* | *D. melanogaster* |
| --- | --- | --- | --- | --- |
| **Total length (bp)** | 220759777 | 231219246 | 193508231 | 143726002 |
| **Scaffolds** | 387 | 338 | 228 | 1870 |
| **N50 (bp)** | 22190323 | 14004682 | 15806012 | 25286936 |
| **L50** | 4 | 5 | 5 | 3 |
| **GC%** | 40.11% | 39.52% | 39.87% | 41.67% |
| **BUSCO Complete, Single Copy** | 93.7% (3078) | 97.8% (3214) | 98.1% (3221) | 98.5% (3235) |
| **BUSCO Complete, Duplicated** | 4.8% (158) | 0.4% (13) | 0.4% (12) | 0.2% (8) |
| **BUSCO Fragmented** | 0.9% (29) | 0.6% (19) | 0.7% (24) | 0.5% (16) |
| **BUSCO Missing** | 0.6% (20) | 1.2% (39) | 0.8% (28) | 0.8% (26) |

### Assembly completeness

BUSCO results for assemblies (Table 1) show a comparable degree of completeness for the 3285 genes in the BUSCO dipteran benchmark set between *D. prolongata* assembly and references, with 3236 complete for *D. prolongata*, 3233 complete for *D. rhopaloa*, and 3243 complete for *D. melanogaster*. The whole genome alignments between the *D. prolongata* assembly and the *D. rhopaloa* (Figure 3A) and *D. melanogaster* references (Figure 3B) further show near complete highly contiguous coverage of the entire reference with regions of *D. prolongata* scaffolds, corresponding to all five major chromosome arms in the *D. melanogaster* genome.

### Repeat annotation

The *D. prolongata* genome exhibits a moderate level of repeat content (24.6%) comparable to the other species (Figure 4). The vast majority (37/40) of classified repeat families are not specific to *D. prolongata*, except for two Long Interspersed Nuclear Element (LINE) retrotransposons, RTE-BovB and L1, and one DNA transposon, Crypton-V (Table S2). We note, however, that further evidence is required to test whether these repeat families have evolved in *D. prolongata*, as all of them have only one identified member in one out of five RepeatModeler runs. Among the repetitive elements of *D. prolongata*, the most prominent repeat classes are Long Terminal Repeats retrotransposons (LTR, 32.2%), LINE (15.1%) and Tandem Repeats (14.6%, Table 2). A breakdown of repeat content by scaffolds across four species can be found in Table S3.

**Figure 4.**
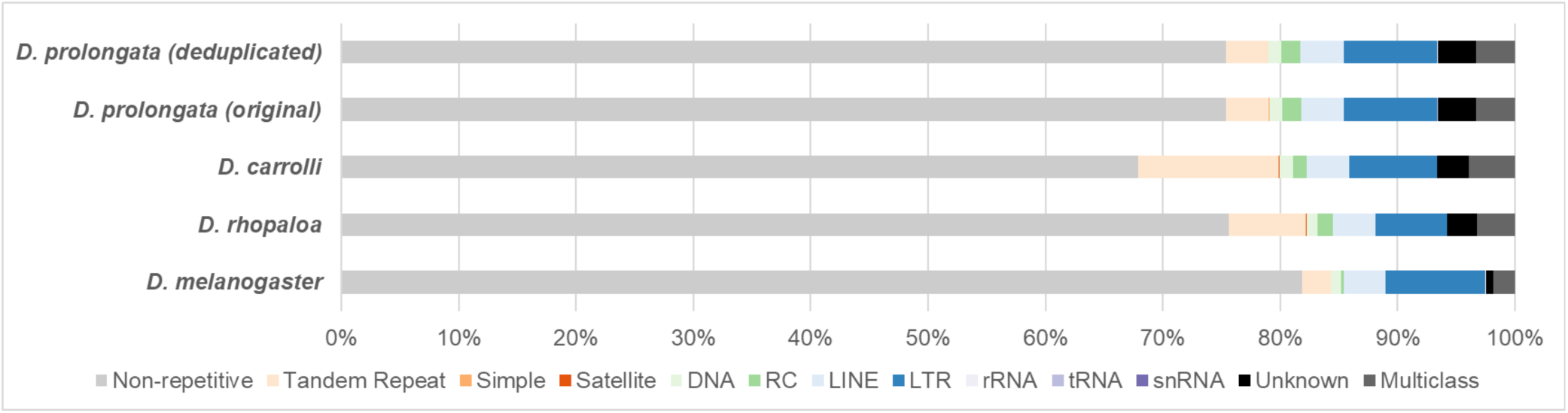
Genome-wide repeat content of *D. prolongata* (before and after de-duplication) and related species. Repeat contents are color coded as follows. Low-complexity regions (Tandem repeats, simple repeats, Satellite): orange palette, DNA transposons (DNA, RC): green palette, retrotransposons (LINE, LTR): blue palette, RNA: purple palette. Abbreviations for each repeat class are as follows. RC: Rolling Circle transposons, LINE: Long-Interspersed Nuclear Element, LTR: Long-Terminal-Repeats retrotransposon, snRNA: small-nuclear RNA, Unknown: unknown class of repeats/transposons, Multiclass: sequences belonging to more than one repeat class.

**Table 2:** Repeat content of genome assemblies of *D. prolongata* and three reference species.

| Repeat Class | *D. prolongata* (%) | *D. carrolli* (%) | *D. rhopaloa* (%) | *D. melanogaster* (%) |
| --- | --- | --- | --- | --- |
| Tandem Repeat | 3.627 | 12.003 | 6.601 | 2.421 |
| Simple | 0.008 | 0.007 | 0.008 | 0.007 |
| Satellite | 0.019 | 0.017 | 0.012 | 0.031 |
| DNA | 1.067 | 1.224 | 0.971 | 0.877 |
| RC | 1.595 | 1.122 | 1.274 | 0.218 |
| LINE | 3.727 | 3.626 | 3.612 | 3.526 |
| LTR | 7.939 | 7.505 | 6.141 | 8.525 |
| rRNA | 0.061 | 0.014 | 0.000 | 0.040 |
| snRNA | 0.000 | 0.001 | 0.004 | 0.000 |
| tRNA | 0.005 | 0.001 | 0.004 | 0.005 |
| Unknown | 3.276 | 2.686 | 2.519 | 0.575 |
| Multiclass | 3.311 | 3.926 | 3.260 | 1.896 |
| **Total** | **24.636** | **32.131** | **24.406** | **18.121** |

Compared with most long (>1Mb) scaffolds, intermediate-sized scaffolds in *D. prolongata* assembly tend to show higher repeat content (Figure S2, Figure S3). Exceptions are found in scaffolds 414, scaffold 293, scaffold 164 and scaffold 280 (Figure S2), where LTR and LINE are overrepresented, reminiscent of the repeat profiles of several primary scaffolds in closely related species *D. carrolli* and *D. rhopaloa* (Figure S4, Figure S5), as well as the Y chromosome in *D. melanogaster* (Figure S6).

### Annotation completeness

Transcripts extracted from the annotation and assembly show that the *D. prolongata* and *D. carrolli* annotations have a high degree of completeness. However, they do not match the completeness of the *D. rhopaloa* and especially *D. melanogaster* references (Table 3), both in terms of gene inclusion and completeness of individual gene models. A higher number of BUSCO dipteran benchmark genes are missing in the *D. prolongata* (95) and *D. carrolli* (115) annotations compared to the *D. rhopaloa* (15) or *D. melanogaster* (0) references. Additionally, the transcripts in the *D. prolongata* and *D. carrolli* annotations are shorter than those from the references, and many more BUSCO dipteran benchmark genes are fragmented in the *D. prolongata* (109) and *D. carrolli* (89) annotations than for *D. rhopaloa* and D. *melanogaster* (both 3). These statistics show the limitations of current algorithmic annotation methods and indicate that care should be used when using gene models from these draft annotations. Despite these limitations, the overall completeness is quite high, with 93.8% of BUSCO benchmark genes covered in both *D. prolongata* and *D. carrolli* annotations, and comparable median transcript lengths in both. These gene models will provide a good foundation for future genetic studies in *D. prolongata* and relatives when used with the limitations of draft annotations in mind. Future iterations of the annotations, when informed by more transcriptome data, will improve gene model coverage and completeness.

**Table 3:**
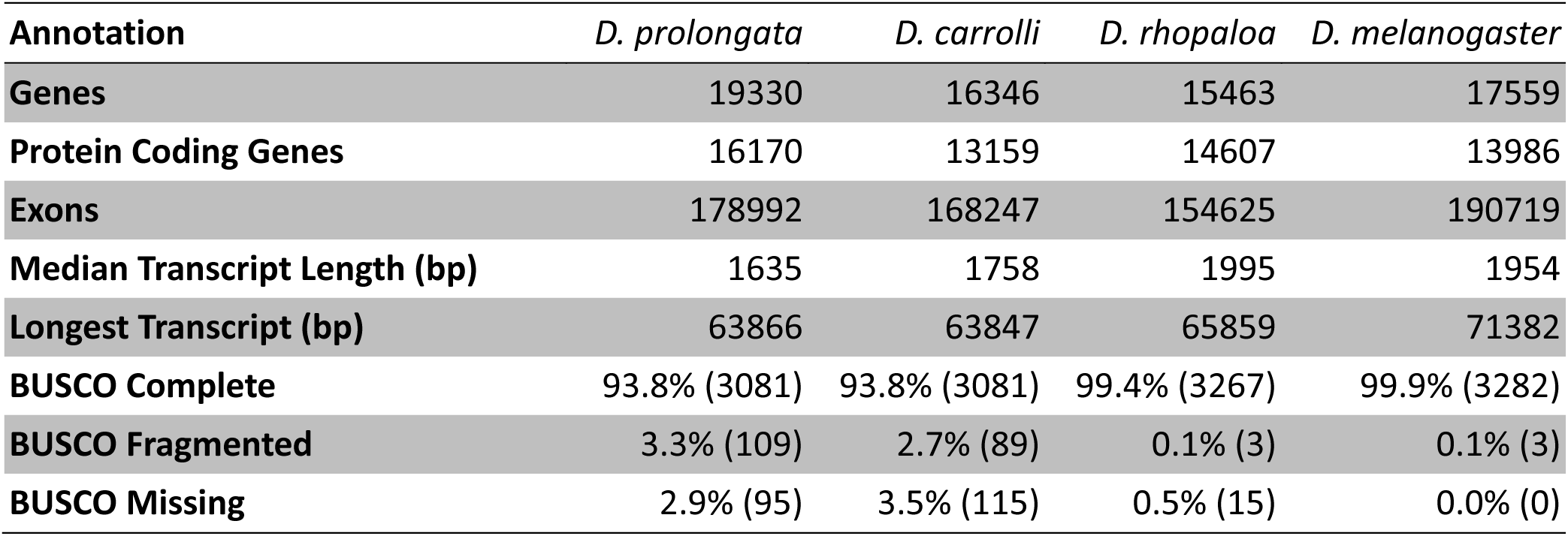
Statistics for annotation completeness for *D. prolongata* and *D. carrolli* annotations alongside reference annotations *D. rhopaloa* GCF_018152115.1 and *D. melanogaster* GCF_000001215.4. BUSCO statistics are for the 3285 genes in the diptera_odb10 benchmark set.

| Annotation | *D. prolongata* | *D. carrolli* | *D. rhopaloa* | *D. melanogaster* |
| --- | --- | --- | --- | --- |
| **Genes** | 19330 | 16346 | 15463 | 17559 |
| **Protein Coding Genes** | 16170 | 13159 | 14607 | 13986 |
| **Exons** | 178992 | 168247 | 154625 | 190719 |
| **Median Transcript Length (bp)** | 1635 | 1758 | 1995 | 1954 |
| **Longest Transcript (bp)** | 63866 | 63847 | 65859 | 71382 |
| **BUSCO Complete** | 93.8% (3081) | 93.8% (3081) | 99.4% (3267) | 99.9% (3282) |
| **BUSCO Fragmented** | 3.3% (109) | 2.7% (89) | 0.1% (3) | 0.1% (3) |
| **BUSCO Missing** | 2.9% (95) | 3.5% (115) | 0.5% (15) | 0.0% (0) |

### Potential regional duplications

The other major caveat for this assembly and annotation is the extent of identified duplication, even after removing duplicate scaffolds. This stands out most clearly in the *D. prolongata* assembly BUSCO scores, where 158 benchmark single-copy genes were identified as duplicated compared to 12 for *D. rhopaloa* and 8 for *D. melanogaster* (Table 1). Additional signals of duplicated regions include the total length of the draft assembly and total gene number in the annotation, which are both higher than in the *D. melanogaster* and *D. rhopaloa* references (Tables 1 and 3), and duplicated regions visible in the whole genome alignment (Figure 3). This suggests some genome regions are represented more than once in the assembly, in addition to any true *D. prolongata*-specific duplication events. Our duplicate gene labeling method identified 945 of 19330 genes (4.89%, close to the BUSCO duplicate frequency); these results are included in Table S4, with a list of duplicated genes on Sheet 1 and the regions and relationships between pairs on Sheet 2; care should be taken when working with these genes and regions. We note that all major (>1Mb) scaffolds in *D. prolongata* have duplicated BUSCO genes even after removal of the fully duplicate scaffolds (Table S1, Figure S3). In contrast, removed scaffolds tend to be intermediate in size and have less repeat content (Figure S7). Remaining BUSCO duplications per scaffold for the final assembly are provided in Sheet 2 of Table S1.

Duplication artifacts often result from heterozygosity persisting through inbreeding (Guo et al. 2016; Kardos et al. 2018; Smith et al. 2019). Segregating inversions, in particular, can capture stretches of heterozygosity and cause the assembler to split haplotypes into separate scaffolds. Consistent with this explanation, the largest remaining duplication candidate visible in the whole genome alignment spans a segregating inversion (Figure 3A’). Sorting biologically real from artifactual duplicates is a key area of improvement for future *D. prolongata* assemblies.

## Data Availability

The final deduplicated assembly for this Whole Genome Shotgun project has been deposited at DDBJ/ENA/GenBank under the accession JAYMZC000000000; the version described in this paper is version JAYMZC010000000. All sequence data used for genome annotation have been deposited in the NCBI Sequence Read Archive under BioProject PRJNA1057277. Genome annotation files for *D. prolongata* and *D. carrolli,* the Dovetail Falcon and HiRise assemblies (containing duplicate scaffolds), sequence file for removed duplicate scaffolds, and all sequence and information files provided by Dovetail have been uploaded to Dryad (URL TBD).

## Acknowledgements

We thank Dr. Hisaki Takamori for providing the original SaPa01 and BaVi44 strain of *D. prolongata*. This work was supported by NIH grant R35 GM122592 to A.K., by NSF award 1601130 to D.L., by the UC Davis Center for Population Biology Pengelley Award to D. L., by UC Davis Center for Population Biology Research Award to Y. L., and by funds from the UNC School of Medicine Strategic Plan to CDJ.

---

**Figure S1.**
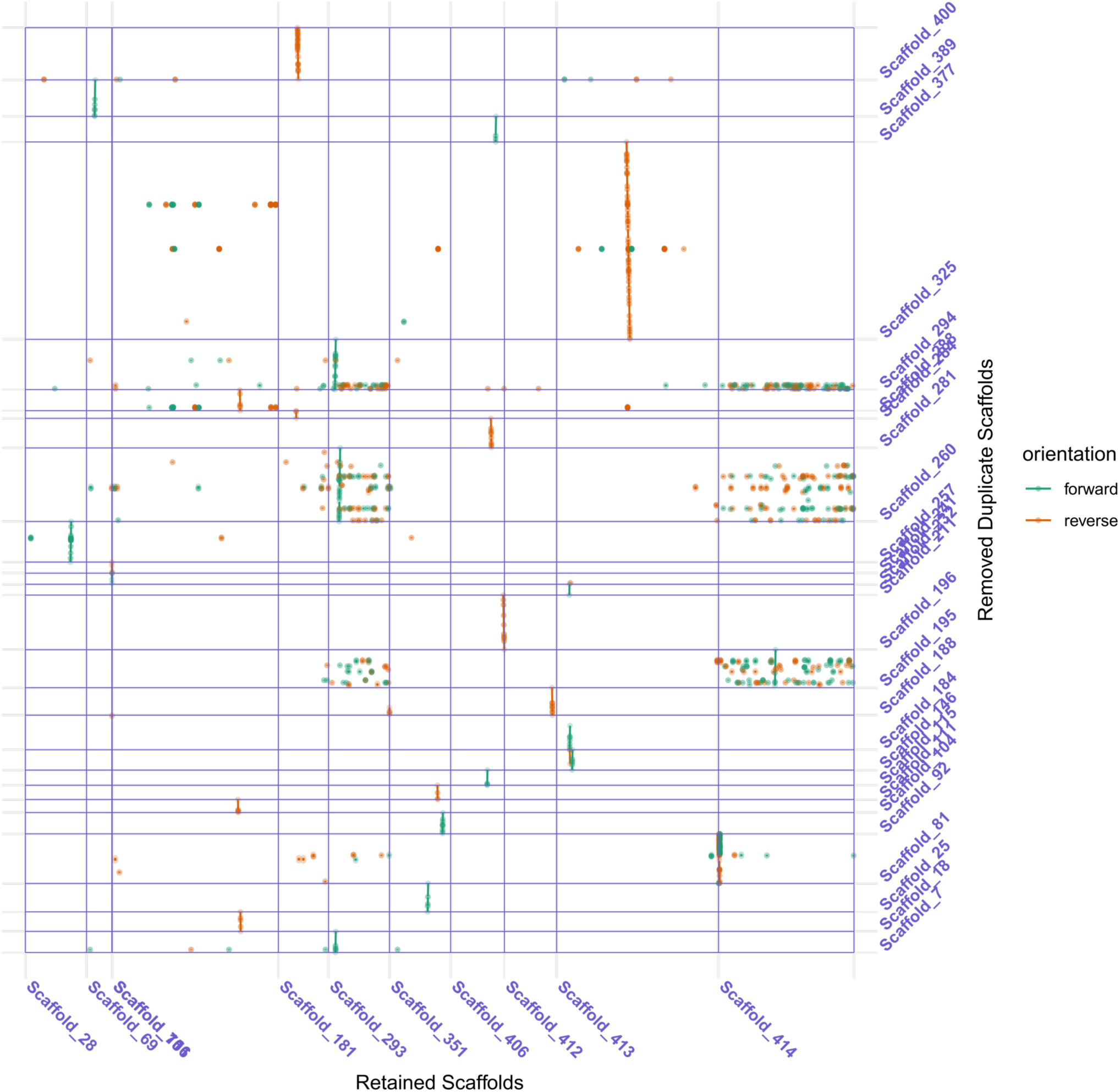
Pairwise MUMmer alignments between 27 duplicate scaffolds and sister scaffolds. Straight lines show alignment between duplicate scaffolds (y-axis) and sister scaffolds (x-axis), with alignment boundaries indicated by flanking points. Sense alignment between scaffolds is shown in green, and antisense alignment is in orange.

**Figure S2.**
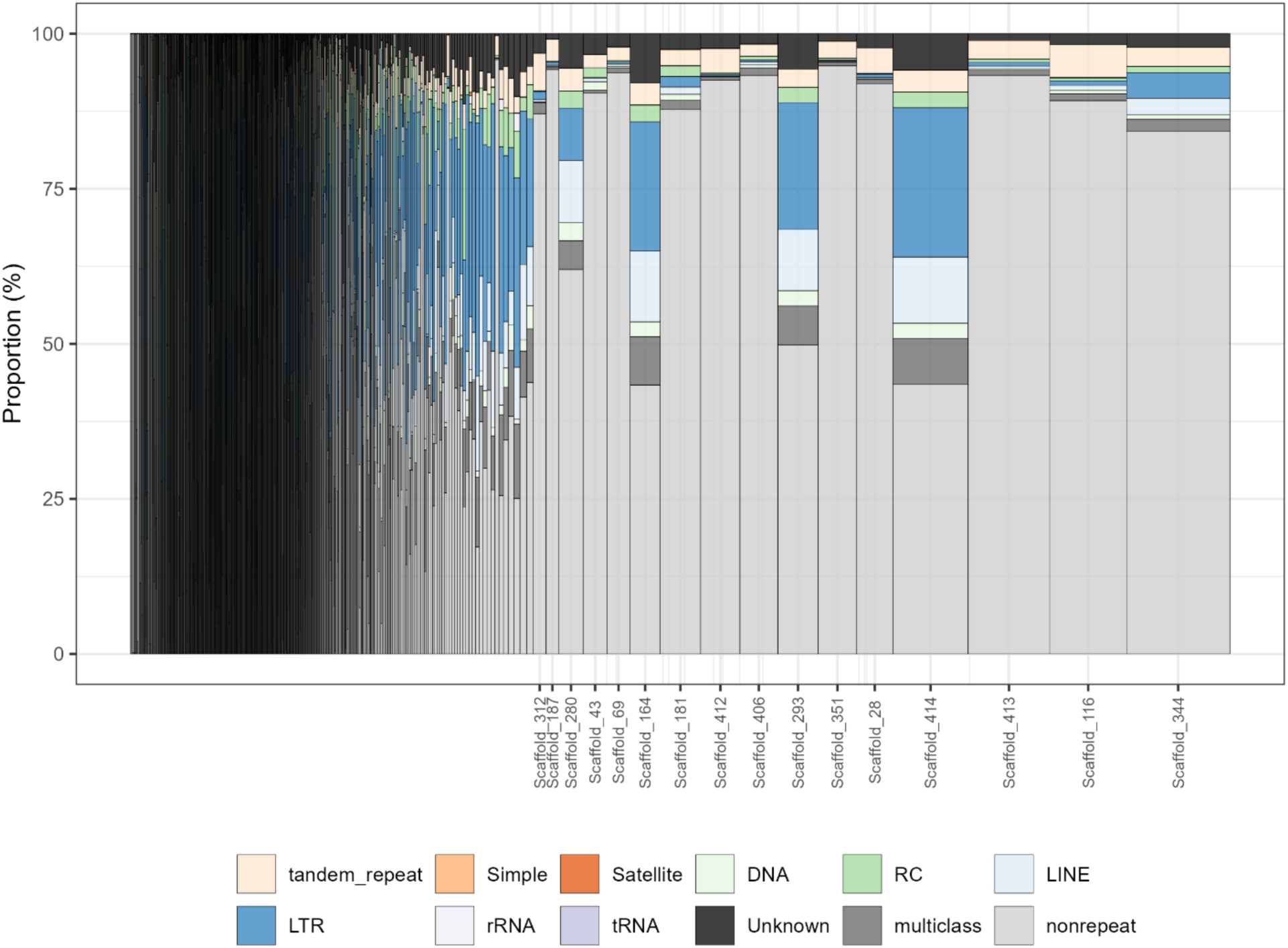
Stacked bar plots showing the distribution of repeat content by scaffolds in *D. prolongata* genome assembly (deduplicated). Widths of bars are proportional to the square root of scaffold/chromosome lengths. Scaffold names are displayed for those of length 1Mb or greater; see Figure S3 for results from smaller scaffolds. Repeat contents are color coded as follows. Low-complexity region: orange palette, DNA transposon: green palette, retrotransposon: blue palette, RNA: purple palette. Abbreviations for each repeat class are as follows. RC: Rolling Circle transposons, LINE: Long-Interspersed Nuclear Element, LTR: Long-Terminal-Repeats retrotransposon, snRNA: small-nuclear RNA, Unknown: unknown class of repeats/transposons, multiclass: sequences belonging to more than one repeat class, nonrepeat: non-repetitive DNA sequence.

**Figure S3.**
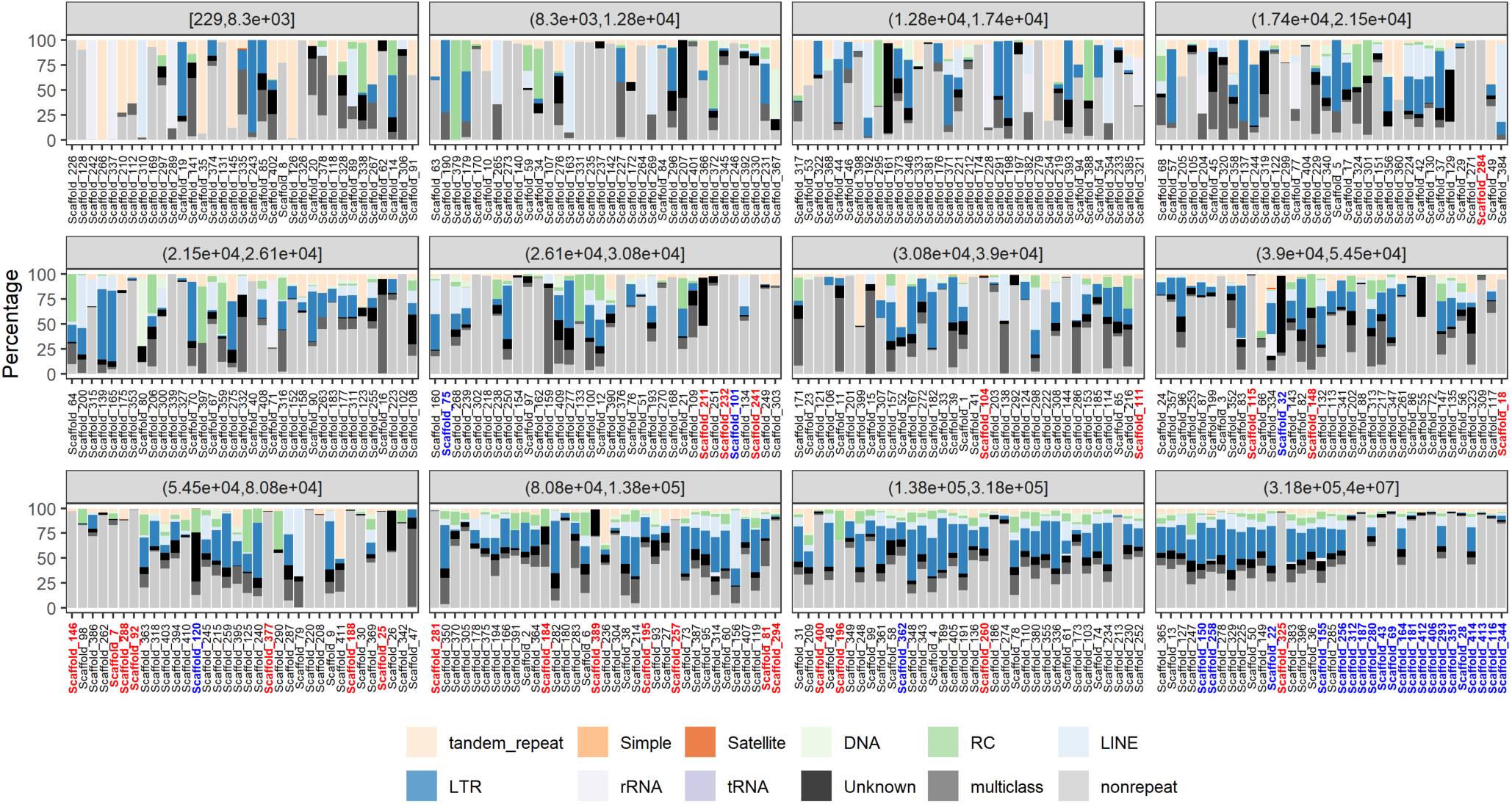
Stacked bar plots showing the distribution of repeat content by scaffolds (partitioned by scaffold length bins) in *D. prolongata* genome assembly. Scaffold names are ordered by their corresponding lengths. Repeat contents are color coded as Fig. S2, with the exception that removed scaffolds have names colored red, and retained members of duplicate scaffold pairs are colored in blue.

**Figure S4.**
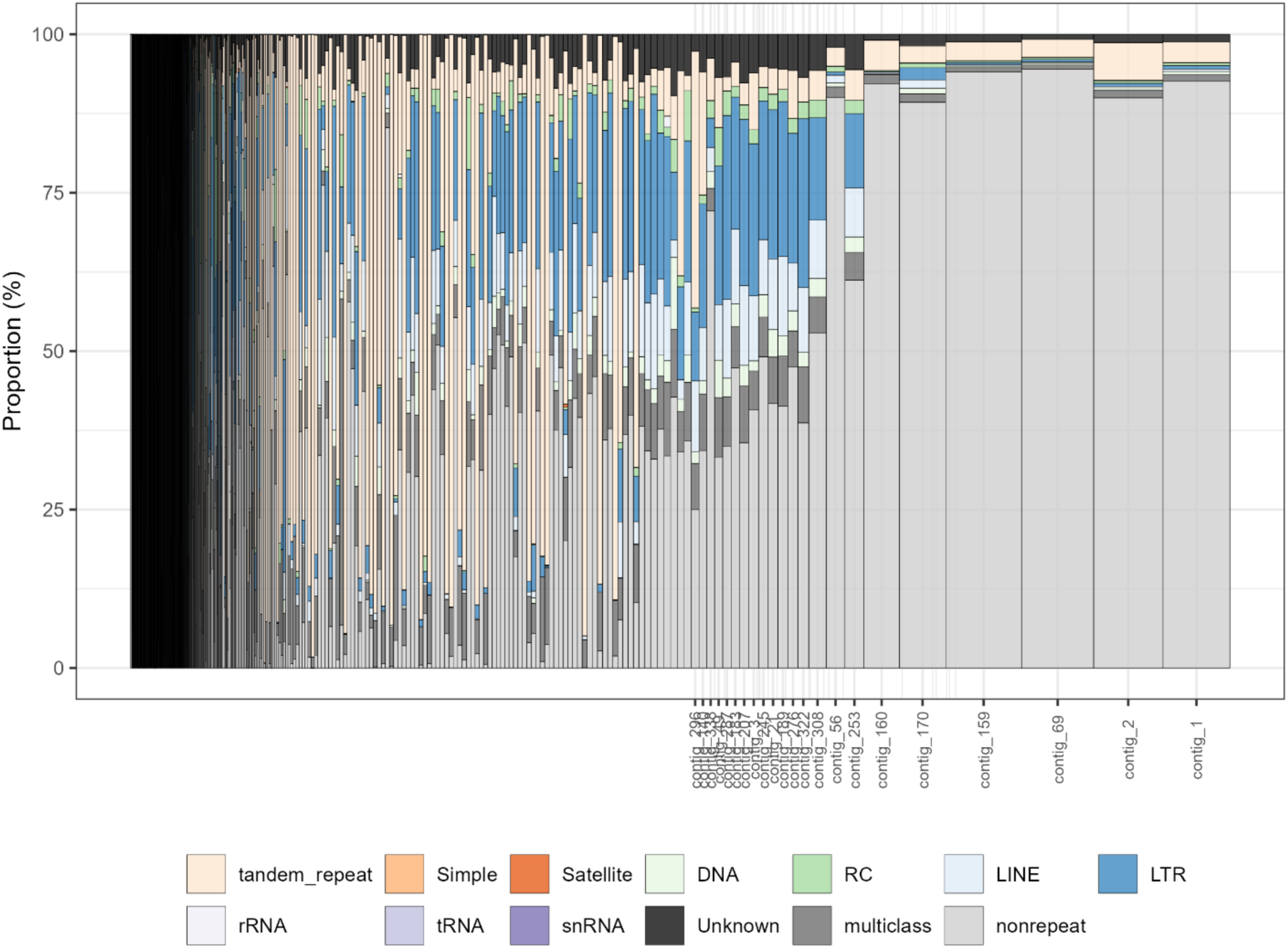
Stacked bar plots showing the distribution of repeat content by scaffolds in *D. carrolli* genome assembly. Widths of bars are proportional to the square root of scaffold/chromosome lengths. Repeat contents are color coded as Fig. S2.

**Figure S5.**
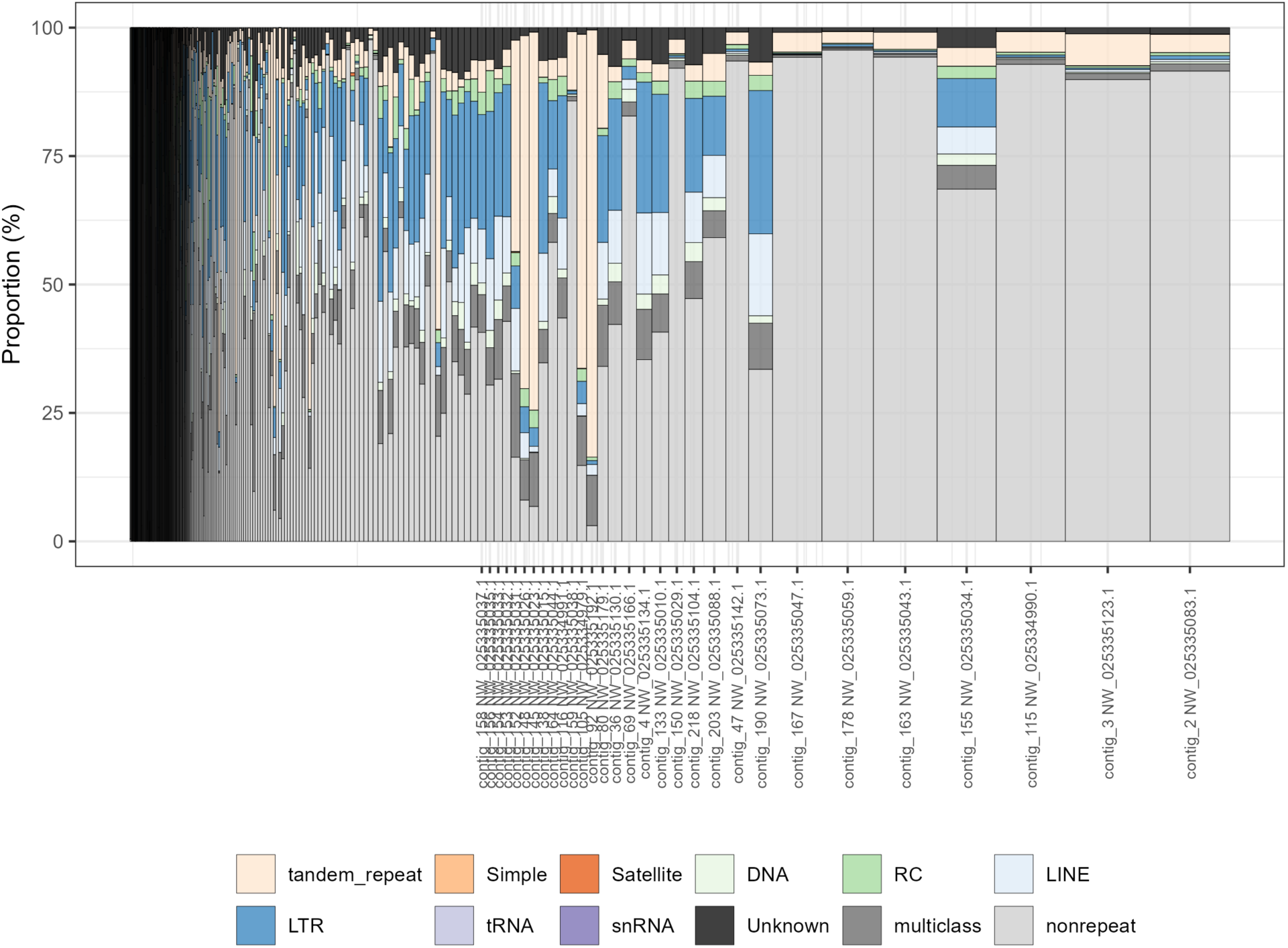
Stacked bar plots showing the distribution of repeat content by scaffolds in *D. rhopaloa* genome assembly (GCF_018152115.1_ASM1815211v1). Widths of bars are proportional to the square root of scaffold/chromosome lengths. Repeat contents are color coded as Fig. S2.

**Figure S6.**
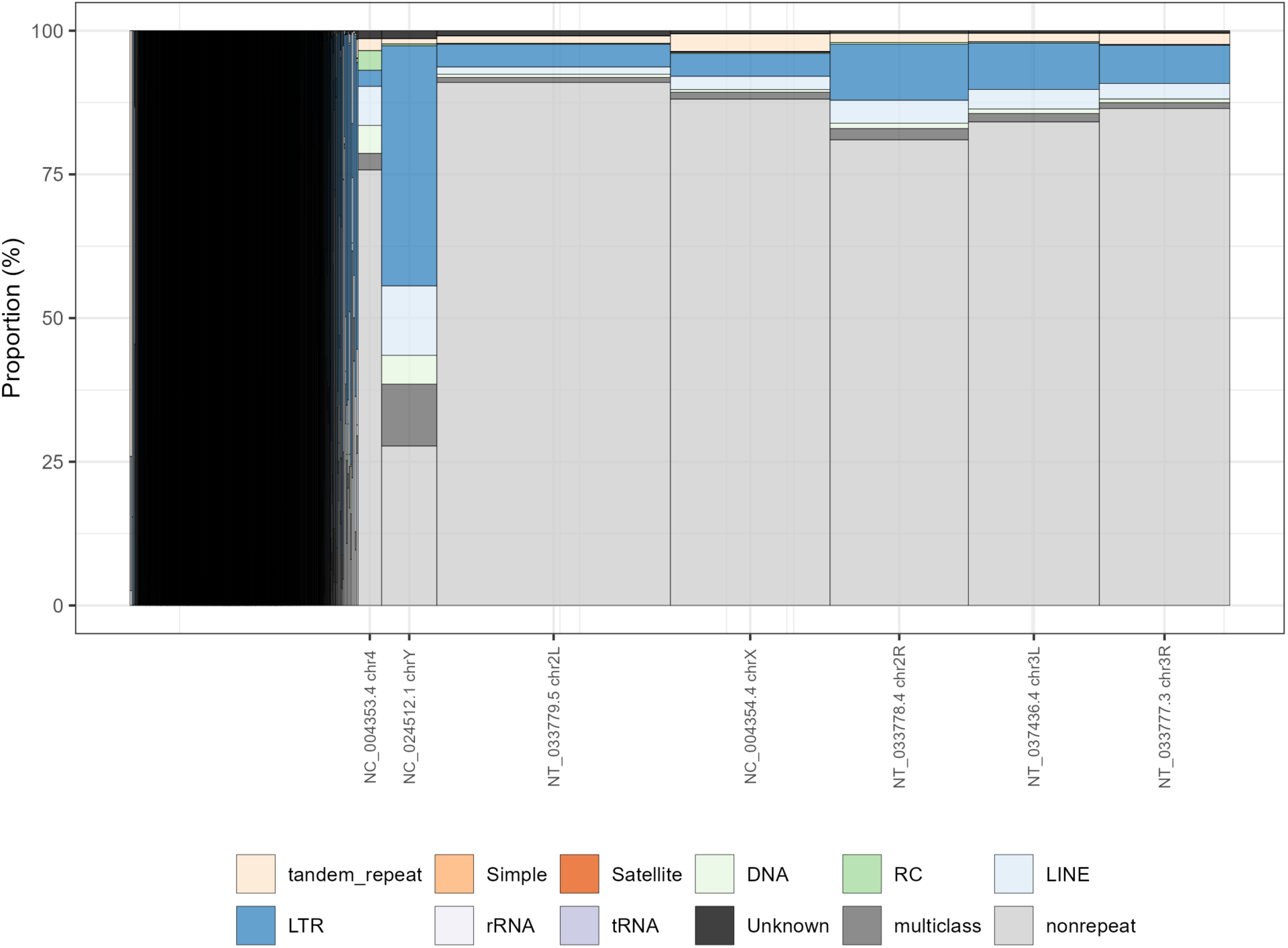
Stacked bar plots showing the distribution of repeat content by scaffolds in *D. melanogaster* genome assembly (GCF_000001215.4_Release_6_plus_ISO1). Widths of bars are proportional to the square root of scaffold/chromosome lengths. Repeat contents are color coded as Fig. S2.

**Figure S7.**
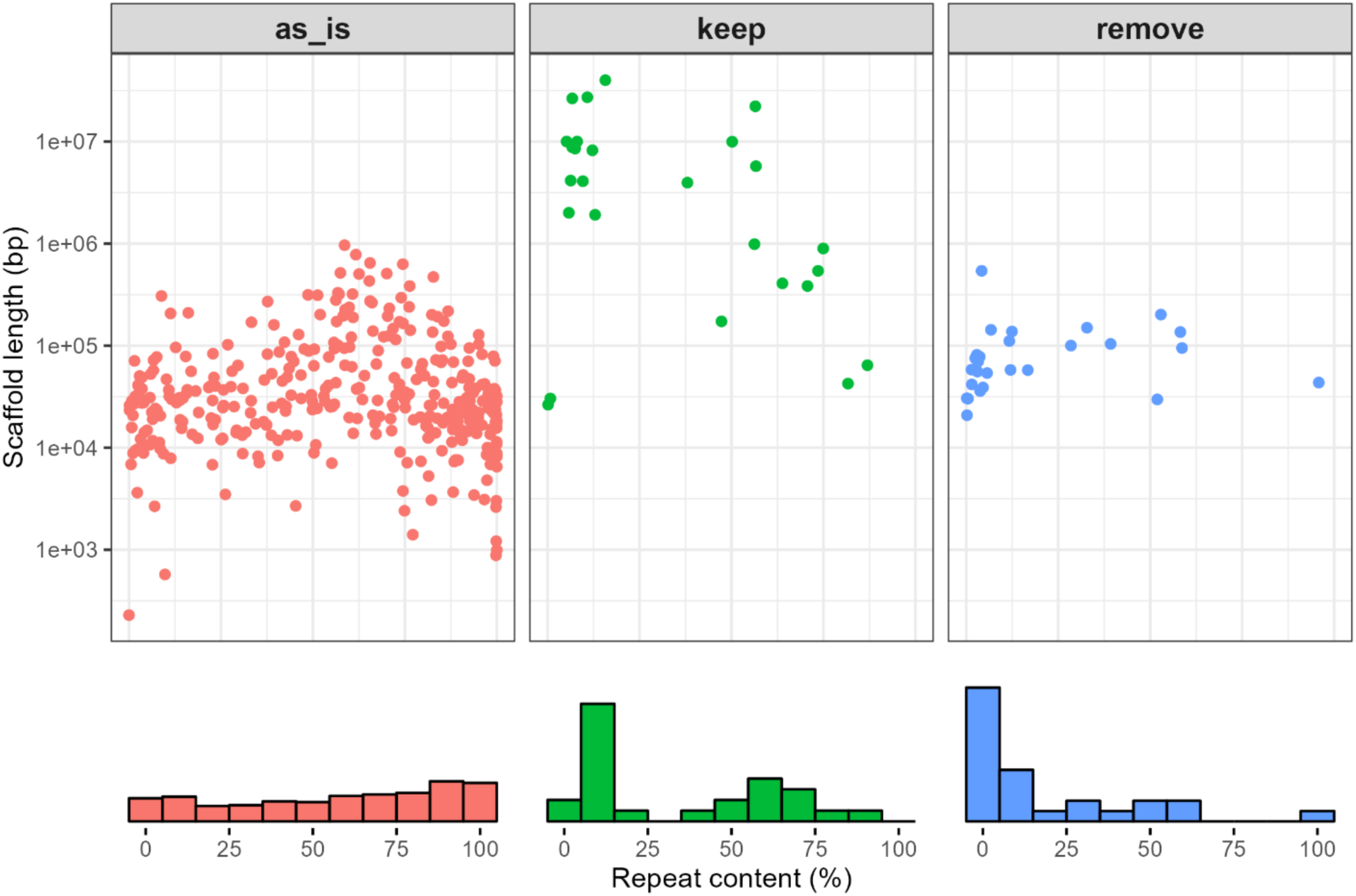
Scatter plots showing the distribution of repeat profiles by scaffolds under each category in the complete *D. prolongata* genome assembly. Frequency histograms of repeat content are displayed at the bottom. X-axis is the repeat content (%), and the y-axis is the corresponding scaffold length. Scaffolds with no BUSCO duplicates are colored in red (as_is), retained scaffolds with BUSCO duplicates in green (keep), and removed scaffolds with BUSCO duplicates in blue (remove).

## Notes

### Competing Interest Statement

The authors have declared no competing interest.

